# Coalescent models at small effective population sizes and population declines are positively misleading

**DOI:** 10.1101/705335

**Authors:** M. Elise Lauterbur

## Abstract

Population genetics employs two major models for conceptualizing genetic relationships among individuals – outcome-driven (coalescent) and process-driven (forward). These models are complementary, but the basic Kingman coalescent and its extensions make fundamental assumptions to allow analytical approximations: a constant effective population size much larger than the sample size. These make the probability of multiple coalescent events per generation negligible. Although these assumptions are often violated in species of conservation concern, conservation genetics often uses coalescent models of effective population sizes and trajectories in endangered species. Despite this, the effect of very small effective population sizes, and their interaction with bottlenecks and sample sizes, on such analyses of genetic diversity remains unexplored. Here, I use simulations to analyze the influence of small effective population size, population decline, and their relationship with sample size, on coalescent-based estimates of genetic diversity. Compared to forward process-based estimates, coalescent models significantly overestimate genetic diversity in oversampled populations with very small effective sizes. When sampled soon after a decline, coalescent models overestimate genetic diversity in small populations regardless of sample size. Such overestimates artificially inflate estimates of both bottleneck and population split times. For conservation applications with small effective population sizes, forward simulations that do not make population size assumptions are computationally tractable and should be considered instead of coalescent-based models. These findings underscore the importance of the theoretical basis of analytical techniques as applied to conservation questions.

## Introduction

Inferring how a population evolves or has evolved under circumstances of interest is a key goal of population genetics. To understand the dynamics of real-world populations, we often turn to analytically and computationally tractable coalescent models of Wright-Fisher population genetics, with the standard being the Kingman coalescent (1982a, 1982b). These models allow the estimation and simulation of current and past population genetic parameters of interest, perhaps foremost among them, genetic diversity or *θ* = 4*N*_*e*_*µ* (Fu and Li 1993; Fu and Li 1999).

Coalescent theory in its standard form (Kingman 1982a, 1982b) both makes certain simplifying assumptions that allow its analytical elegance, and remains the typical model used to interpret genetic variation. These include a stable effective population size large enough to be effectively infinite, that is much larger than the sample size. This is because of the fundamental simplifying assumption that only two lineages coalesce per generation. However when the effective population size is very small (*N*_*e*_ < 1000), and/or the sample size approaches or exceeds the effective population size, the probability of more than two lineages coalescing in a single generation (multiple mergers) increases exponentially (figure 1) (Wakeley 2016).

**Figure 1.**
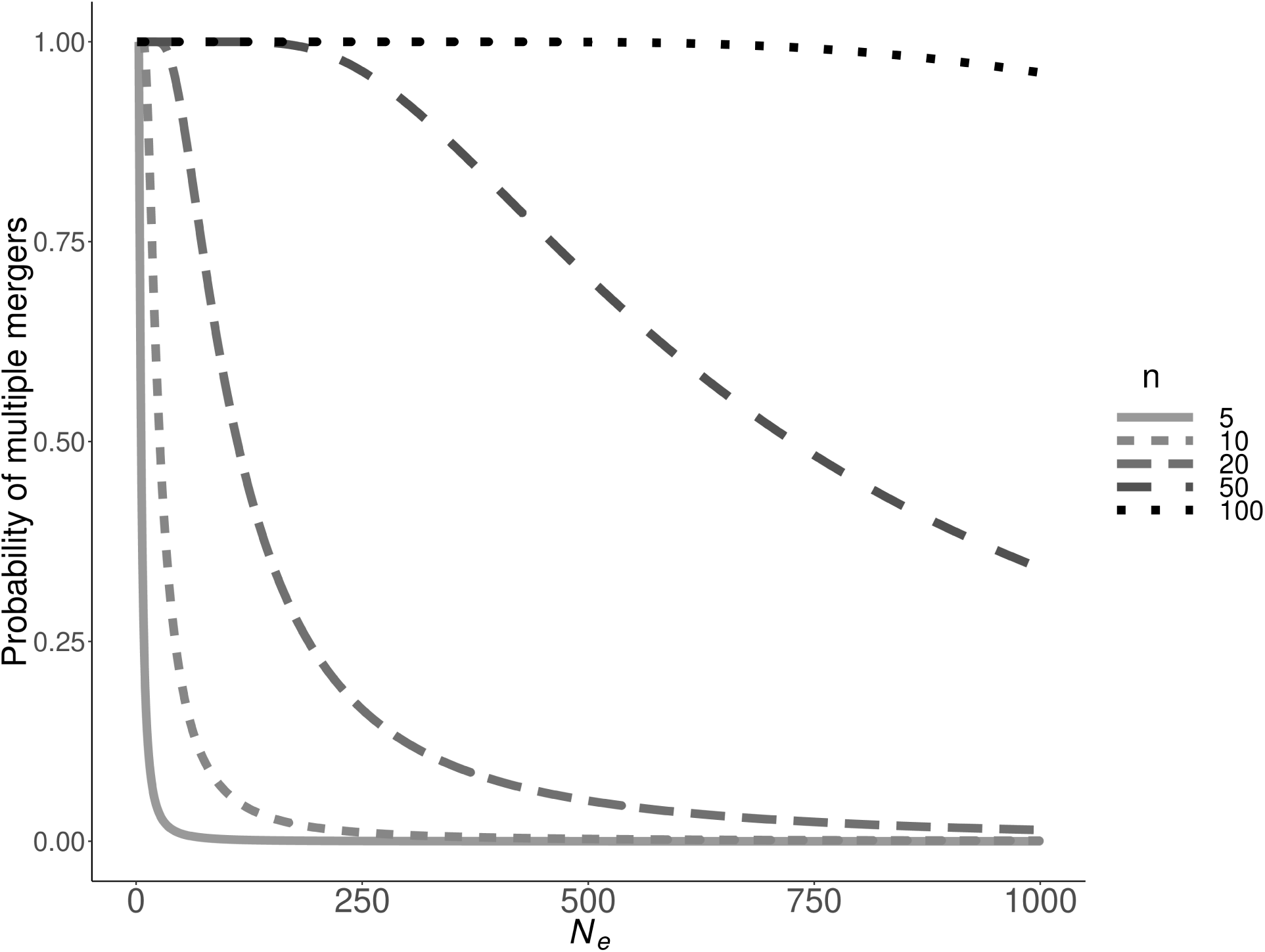
Effective population size and multiple merger probability. The probability that more than two lineages coalesce per generation (multiple mergers) as a function of sample size (n) and effective population size from *N*_*e*_ = 0 to 1000.

In species of conservation concern, as well as widely-sampled species such as humans, one or more of these assumptions (large *N*_*e*_, *N*_*e*_ >> *n*, stable *N*_*e*_) are often unmet. For example, in a sample of 138 studies of species with genetically estimated effective population sizes, Palstra and Ruzzante (2008) found 120 species to have estimated effective population sizes of less than 1000, and 27 to have effective population sizes of less than 100. Included among these are a population North Sea cod (*Gadus morhua*) off the Yorkshire coast of England, with an effective population size of 70 - 120 depending on sampling period (Hutchinson et al. 2003); Brazilian water hyacinth (*Eichhornia paniculata*) in northeastern Brazil, with an effective population size of < 5 - 70 depending on the population sampled (Husband and Barrett 1992); and the Siberian tiger (*Panthera tigris altaica*) with an effective population size of approximately 14 (Alasaad et al. 2011).

Given these small N_e_, some species and populations may be oversampled relative to effective population size. For example, the hihi (*Notiomysts cincta*) of New Zealand was estimated to have an effective population size of 10 and a census size of 30 after reintroduction to Mokoia island (Castro et al. 2004). In this case, because of banding and monitoring, it was possible to sample the entire population, and thus have a sample size three times that of the effective population size. Estimates of human effective population sizes vary depending on population(s) and method of estimation, but are generally in the range of a few thousand to ∼15,000 (Tenesa et al. 2007; Mezzavilla 2015; The 1000 Genomes Project Consortium 2015). This means that current human research involving samples of tens or even hundreds of thousands of individuals (eg. Mazet et al. 2016; Martin et al. 2017) is also likely to be oversampled relative to effective population size.

As in humans, small effective population sizes are often the result of population bottlenecks. Bottlenecks are characteristic of populations of conservation concern as well as the founding of new populations. Immediately after a bottleneck, genetic diversity, as measured by descriptive statistics involving the number of alleles, such as Watterson’s theta (*θ*_*W*_), drops precipitously (Maruyama and Fuerst 1985; Leberg 1992). This is because of the immediate loss of rare alleles, particularly singletons. In contrast, heterozygosity is reduced less immediately after a bottleneck, but continues to fall if the bottleneck is prolonged (Allendorf 1986).

The robustness of coalescent theory to violations of theoretical assumptions has been relatively well-studied with respect to sample size, migration, population size fluctuations, and variation in reproductive success, among others. For example, Wakeley and Takahashi (2003) and Bhaskar et al. (2014) found that large sample sizes, relative to the effective population size, increase the proportion of singleton polymorphisms relative to standard coalescent expectations. Bhaskar et al. (2014) were able to attribute this directly to multiple mergers in oversampled genealogies. In addition, variance in reproductive success can also result in multiple coalescent events per generation, and increases the complexity of the relationship between effective population size and the per-locus neutral mutation rate (Eldon and Wakeley 2005). Migration and recolonization in metapopulations also causes skew in the site frequency spectrum (Wakeley and Aliacar 2001). In these cases, deviations are attributable to multiple mergers.

Intuition might suggest that very small effective population sizes could have the same effects on parameter inference as do other violations resulting in multiple mergers, but the exact nature of the consequences and how they interact with population bottlenecks remains unexplored. As previously suggested, application of multiple merger coalescent models could resolve erroneous estimates of population genetic parameters in populations that violate assumptions of the standard coalescent (Schweinsberg 2000; Mohle and Sagitov 2003; Eldon and Wakeley 2005; Birkner and Blath 2008; Tellier and Lemaire 2014; Koskela, Jenkins, and Spano 2015; Montano 2016). However, if and how this applies to populations with very small effective population sizes (*N*_*e*_ < 1000) and bottlenecks has so far been neglected.

Here, I use simulations to examine the effect of small effective population sizes and population declines, and their interaction with sample size, on coalescent estimates of genetic diversity. I compare coalescent estimates of genetic diversity to genetic diversity generated by forward simulations for a range of constant effective population sizes between *N*_*e*_ = 10 and 1000, and a range of sample sizes between *n* = 2 and 1000. This includes scenarios of oversampling: parameter spaces in which the sample size approaches and even exceeds the effective population size (*n* > *N*_*e*_). Because small effective population sizes are often the result of more or less recent bottlenecks, I extend the simulations to include bottlenecks of severity ranging from 0% to 99% and sampling times (without recovery) between *T* = 1 and 1000 generations post-bottleneck. These comparisons show that coalescent models tend to incorrectly estimate genetic diversity when effective population sizes are very small and when sample size exceeds effective population size. These effects are especially pronounced after a recent bottleneck.

## Materials and Methods

To ascertain the reliability of coalescent-based estimates of genetic diversity at low effective population sizes (*N*_*e*_ < 1000) and sample sizes (n) approaching *N*_*e*_, both with and without bottleneck at varying times (T), I used forward, process-based simulations to ground-truth coalescent simulations (table 1). Unlike coalescent simulations, which probabilistically model the history of a sample backward in time, Wright-Fisher forward simulations follow the entire population of individuals generation by generation from the beginning of the simulation until it ends and individuals are sampled (eg. Hernandez 2008; Haller and Messer 2017, 2019).

**Table 1.** Explored parameter space of *N*_*e*_ and n, with approximate relationship between *N*_*e*_ and n in the cells.

| $n \mid N_e$ | 10 | 20 | 50 | 100 | 200 | 500 | 1000 |
| --- | --- | --- | --- | --- | --- | --- | --- |
| 2 | $n < N_e$ | $n \ll N_e$ | $n \ll N_e$ | $n \ll N_e$ | $n \ll N_e$ | $n \ll N_e$ | $n \ll N_e$ |
| 5 | $n < N_e$ | $n < N_e$ | $n \ll N_e$ | $n \ll N_e$ | $n \ll N_e$ | $n \ll N_e$ | $n \ll N_e$ |
| 10 | $n = N_e$ | $n < N_e$ | $n < N_e$ | $n \ll N_e$ | $n \ll N_e$ | $n \ll N_e$ | $n \ll N_e$ |
| 20 | $n > N_e$ | $n = N_e$ | $n < N_e$ | $n < N_e$ | $n \ll N_e$ | $n \ll N_e$ | $n \ll N_e$ |
| 50 | $n > N_e$ | $n > N_e$ | $n = N_e$ | $n < N_e$ | $n < N_e$ | $n \ll N_e$ | $n \ll N_e$ |
| 100 | $n \gg N_e$ | $n > N_e$ | $n > N_e$ | $n = N_e$ | $n < N_e$ | $n < N_e$ | $n \ll N_e$ |
| 200 | $n \gg N_e$ | $n \gg N_e$ | $n > N_e$ | $n > N_e$ | $n = N_e$ | $n < N_e$ | $n < N_e$ |
| 500 | $n \gg N_e$ | $n \gg N_e$ | $n \gg N_e$ | $n > N_e$ | $n > N_e$ | $n = N_e$ | $n < N_e$ |
| 1000 | $n \gg N_e$ | $n \gg N_e$ | $n \gg N_e$ | $n \gg N_e$ | $n > N_e$ | $n > N_e$ | $n = N_e$ |

### Simulations

Forward simulations were performed using SFS_CODE (Hernandez 2008) and SLiM v2 (Haller and Messer 2017), and coalescent simulations were performed with Hudson’s ms (Hudson 2002). I performed 10,000 simulations of each parameter combination {*N*_*e*_, *n*, T} with SFS_CODE and Hudson’s ms. SLiM v2 was used to verify the SFS_CODE results by performing 10,000 simulations of each parameter combination {*N*_*e*_, *n*} to mutation-drift equilibrium. Effective population size *N*_*e*_ could take the values {10, 20, 50, 100, 200, 500, 1000}, and sample size n the values {2, 5, 10, 20, 50, 100, 200, 500, 1000}, chosen to reflect plausible effective population sizes of populations of conservation concern. For the forward simulations, when sample size exceeded effective population size, individuals were resampled until the sample size was reached. The non-bottleneck simulations were run with constant *N*_*e*_ = {10, 20, 50, 100, 200, 500, 1000}, and forward simulations were run with a burn-in of 10 * 2*N*_*e*_.

For the bottleneck simulations, pre-bottleneck *N*_*e*_ = 1000, since preliminary simulations showed no significant differences between forward and coalescent diversity estimates at *N*_*e*_ = 1000, and because forward simulations were computationally impractical at *N*_*e*_ > 1000. The bottleneck reduced the population to one of the above set of possible *N*_*e*_ values at *T* = {1, 10, 100, 1000} generations, resulting in sampling times ranging from 1 to 1000 generations after the bottleneck event. There was no post-bottleneck recovery of *N*_*e*_. For both non-bottleneck and bottleneck scenarios, a 5,000-base-pair locus was simulated with mutation rate = 4*N*_*e*_ * 10^-6^ and recombination rate = 4*N*_*e*_ * 10^-8^ per bp. See table 1 for {*N*_*e*_, *n*} parameter space.

Since rare variants are lost first during a bottleneck, which reduces the number of segregating sites, Watterson’s *θ* (*θ*_*W*_) is an appropriate and sensitive summary statistic for comparing genetic diversity estimated by each type of simulation (Allendorf 1986). It is also commonly used in studies of genetic diversity both within and among populations. I compared *θ*_*W*_ estimated by each type of simulation at each parameter combination {*N*_*e*_, *n, T*}.

### Statistical comparisons

Each set of 10,000 simulations per parameter combination {*N*_*e*_, *n, T*} and simulation type (forward, coalescent) created a distribution of estimated *θ*_*W*_ values. At small *N*_*e*_, the distributions of the number of segregating sites, thus *θ*_*W*_, is zero-inflated, and parametric tests are inappropriate. To compare mean *θ*_*W*_ among coalescent and both forward simulators for each set of simulations with the same parameter combination (without a bottleneck), I used Kruskal-Wallis tests followed by a post-hoc Dunn’s test for each significant case. For the bottleneck scenarios, I used Kolmogorov-Smirnov tests to compare the distributions, and Mann-Whitney U tests to determine if the coalescent and forward samples could have come from a distribution with the same mean *θ*_*W*_, between pairs of forward-coalescent distributions created with the same parameter combination. All p-values were converted to false discovery rate (FDR) with a threshold of 0.001 to control for multiple comparisons.

### Data Availability

Supplementary tables have been uploaded to figshare, and code and data for analyses have been uploaded to https://github.com/lauterbur/Ne.

## Results

### Constant small effective population size

There were few differences between genetic diversity (as measured by Watterson’s *θ*) calculated from both forward simulators, SFS_CODE and SLiM v2. The post-hoc Dunn’s test showed false discovery rates (FDR) under the 0.001 threshold only occurred for 4 out of 63 comparisons (6%). Thus, only comparisons between SFS_CODE (forward) and Hudson’s ms (coalescent) are shown and used for subsequent bottleneck analyses (Table S1).

For those scenarios in which significant differences between *θ*_*W*_ calculated from SFS_CODE and SLiM simulations were found, these differences were 16-30% the magnitude of differences between the coalescent estimate and either forward *θ*_*W*_. All such scenarios had sample sizes approximately double the effective population size ({*N*_*e*_ = 100, *n* = 200}, {*N*_*e*_ = 200, *n* = 500}, {*N*_*e*_ = 500, *n* = 1000}). Hence, the discrepancy could be an effect of the resampling of individuals performed when sample size exceeds effective population size.

At constant effective population size, statistically significant differences between *θ*_*W*_ estimated by coalescent models and calculated from forward models were most pronounced when sample size was much different from effective population size (Kruskal Wallis FDR < 0.001 and Dunn’s FDR < 0.001). When *N*_*e*_ ≥ *n*, the coalescent overestimated genetic diversity, and when *N*_*e*_ < *n*, the coalescent underestimated genetic diversity (figure 3).

**Figure 2.**
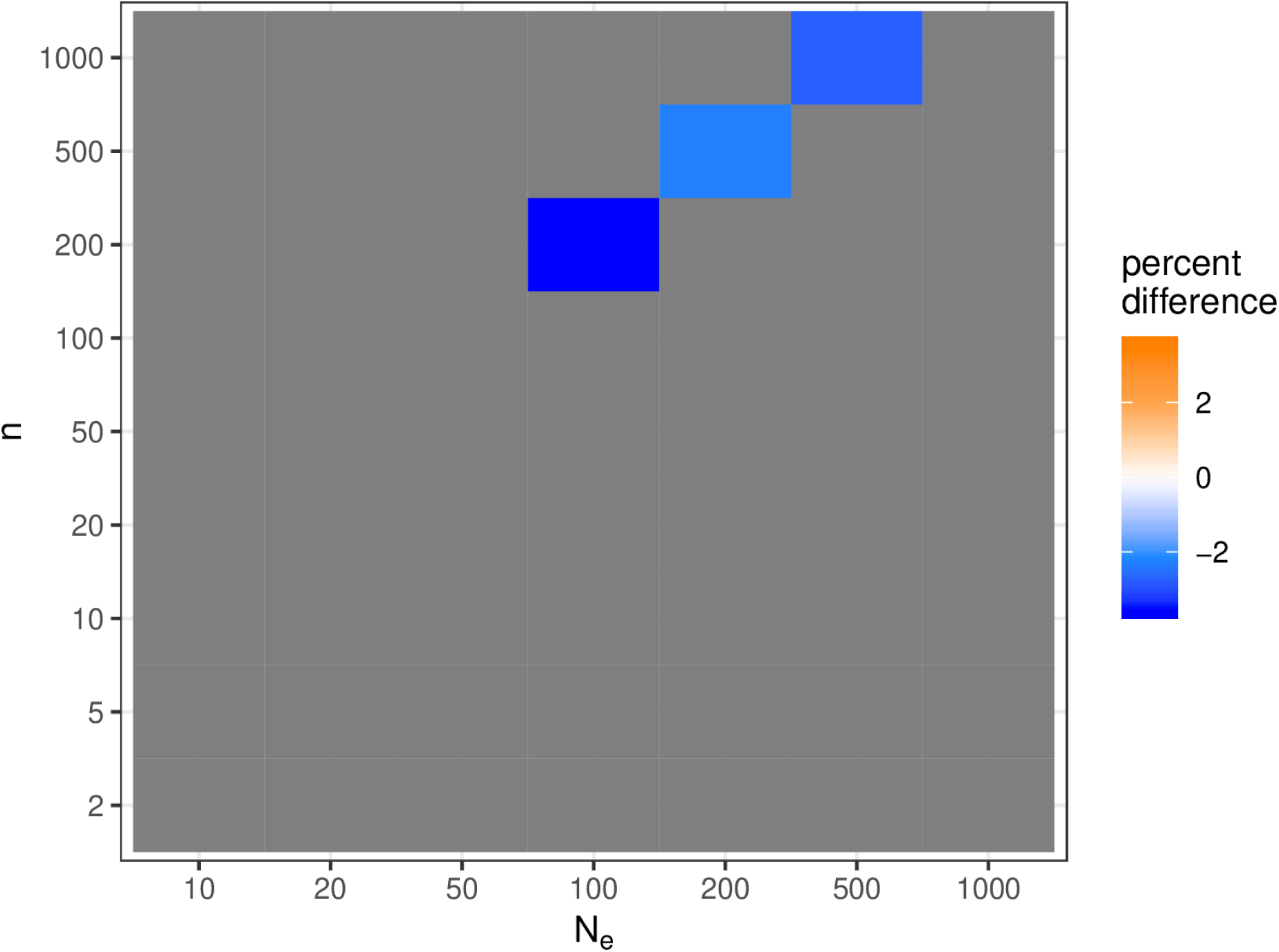
Significant differences between *θ*_*W*_ values for SLiM and SFS_CODE. Grey indicates no significant difference, colors indicate significant differences with darker oranges associated with larger mean percent differences.

**Figure 3.**
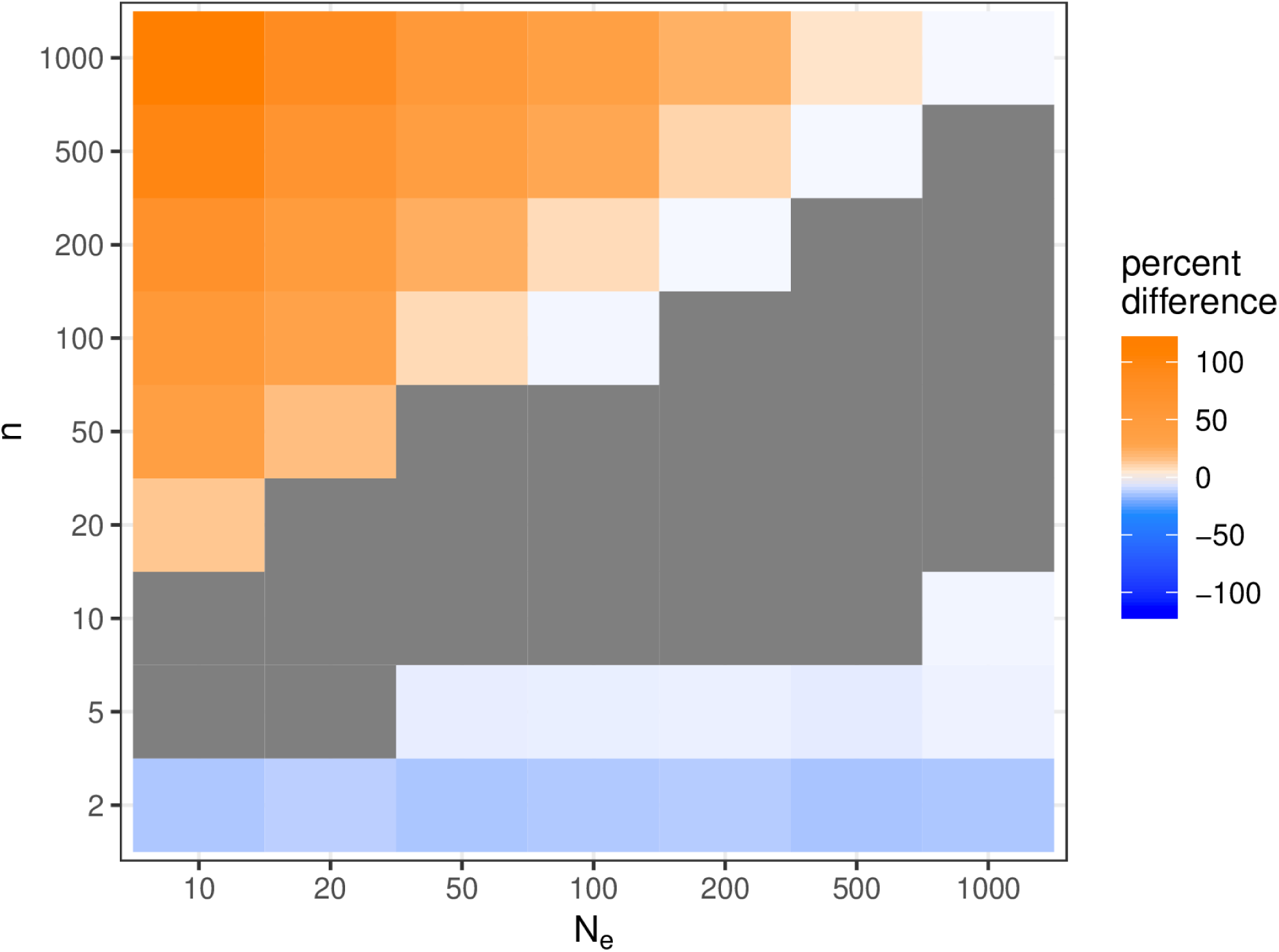
Significant differences between *θ*_*W*_ values for forward (SFS_CODE) and coalescent (ms) simulations. Grey indicates no significant difference, colors indicate significant differences with darker oranges associated with larger mean positive percent differences and darker blues associated with larger mean negative percent differences.

The relationship between sample size (*n*) and effective population size (*N*_*e*_) influenced the result of forward to coalescent comparisons of genetic diversity. At small *n/N*_*e*_ ratios (≤ 0.1), coalescent simulations were more likely to underestimate *θ*_*W*_ by up to 13%, especially at small *N*_*e*_, while at large *n/N*_*e*_ ratios (≥ 2.0) coalescent simulations were more likely to overestimate *θ*_*W*_ by up to 60% (figure 4)

**Figure 4.**
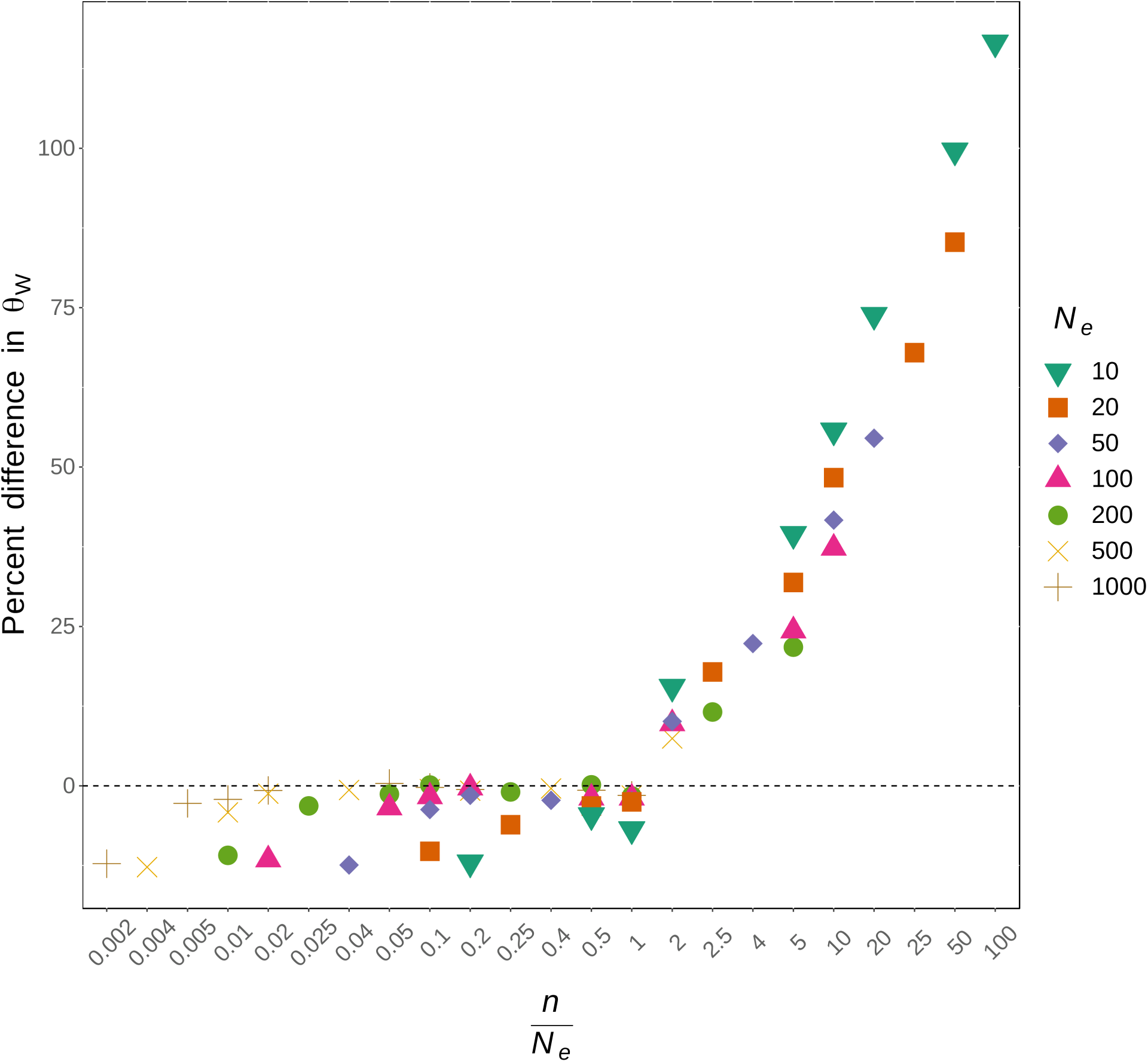
The effect of the relationship between sample size and effective population size (*n/N*_*e*_) on coalescent estimates of *θ*_*W*_ relative to forward calculations of *θ*_*W*_. When the percent difference is positive, the coalescent estimate is larger and thus overestimating *θ*_*W*_, and when the percent difference is negative, the coalescent estimate is smaller and thus underestimating *θ*_*W*_.

### Bottlenecks to small effective population size

The time between sampling and a population bottleneck in small populations had a greater effect on the accuracy of coalescent estimates of *θ*_*W*_ than effective population size and its relationship with sample size in populations of constant size (Table S2). Immediately after a bottleneck (*T* = {1, 10}), coalescent estimates of *θ*_*W*_ overestimate genetic diversity at all effective population sizes except *N*_e_ = 1000, though not all comparisons at *N*_e_ = 200 and *N*_e_ = 500 are significant. At *N*_e_ = 1000, coalescent estimates underestimate genetic diversity (figure 5, figure 6). There was no effect of the relationship between sample size and effective population size at *T* = 1. At *T* = 10 there was an effect only at the smallest effective population sizes (*N*_*e*_ = {10, 20}, figure 7a, b).

**Figure 5.**
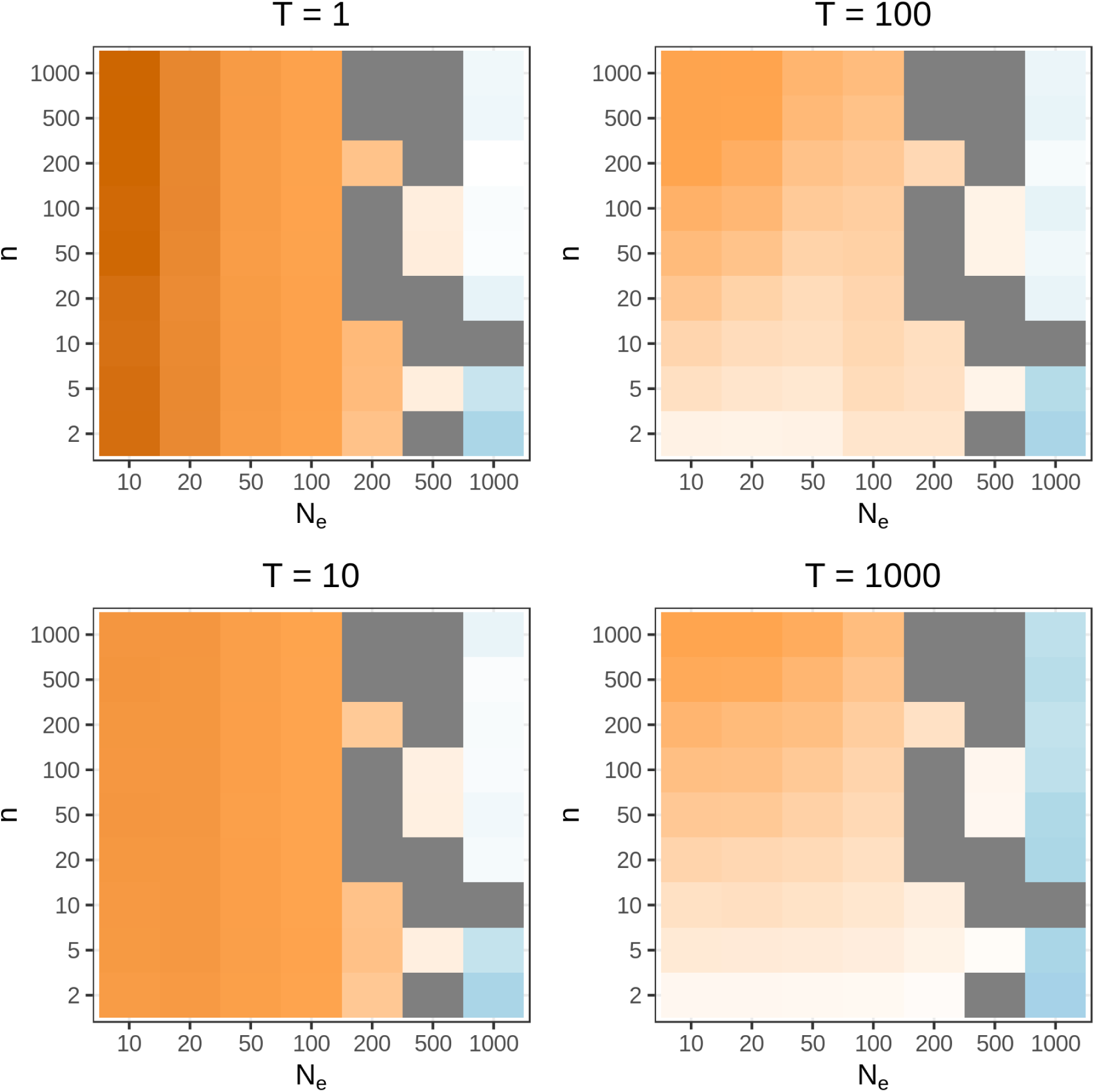
Significant differences between *θ*_*W*_ values for forward (SFS_CODE) and coalescent (ms) simulations, by time since bottleneck (T). Grey indicates no significant difference, colors indicate significant differences with darker oranges associated with larger mean positive percent differences and darker blues associated with larger mean negative percent differences.

**Figure 6.**
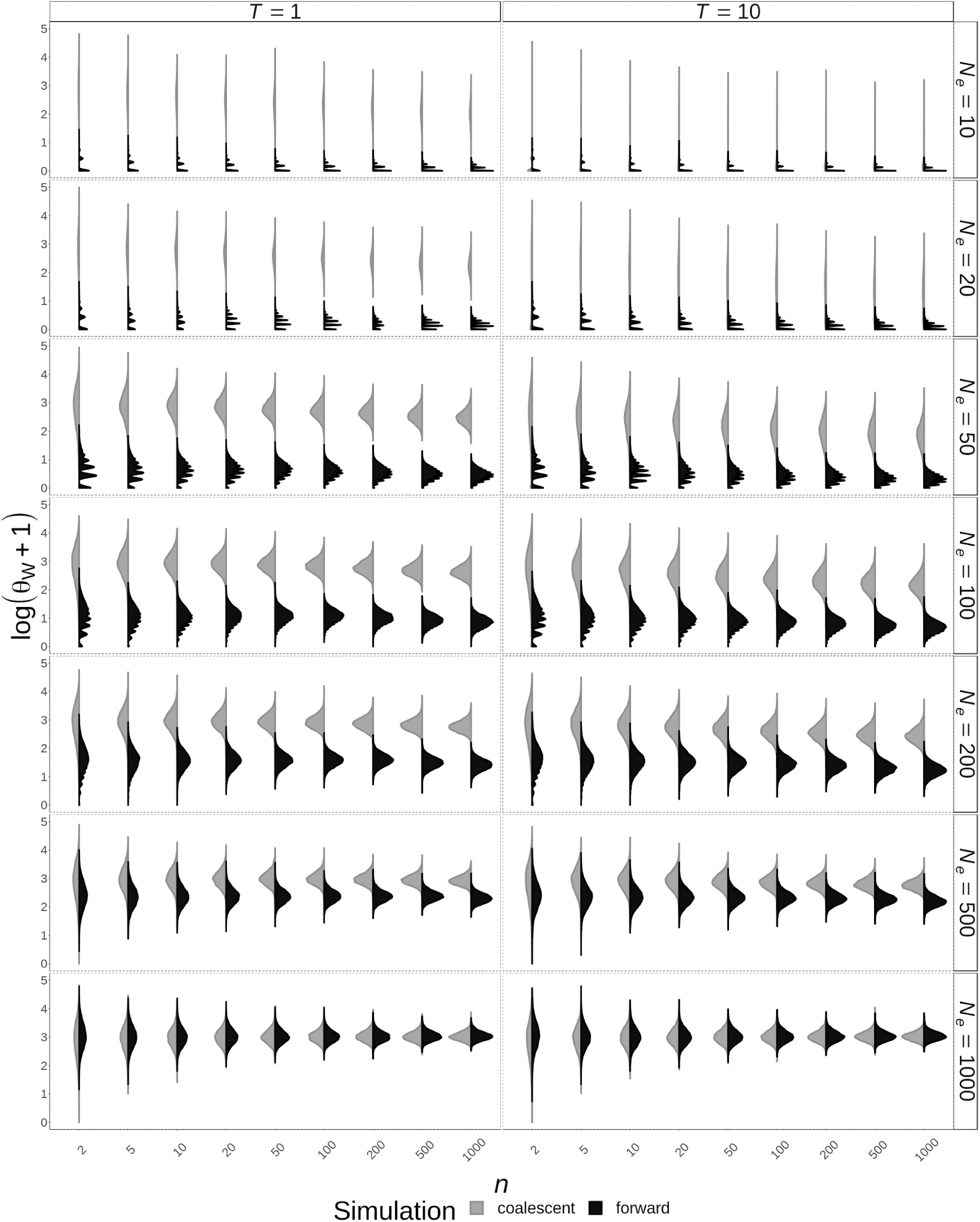
Distributions of *θ*_*W*_ (log transformed for visibility) estimated from coalescent simulations (grey) compared to *θ*_*W*_ calculated from forward simulations (black), across sample size (x axis) and effective population size (vertical facets) for *T* = 1 (left) and *T* = 10 (right).

**Figure 7.**
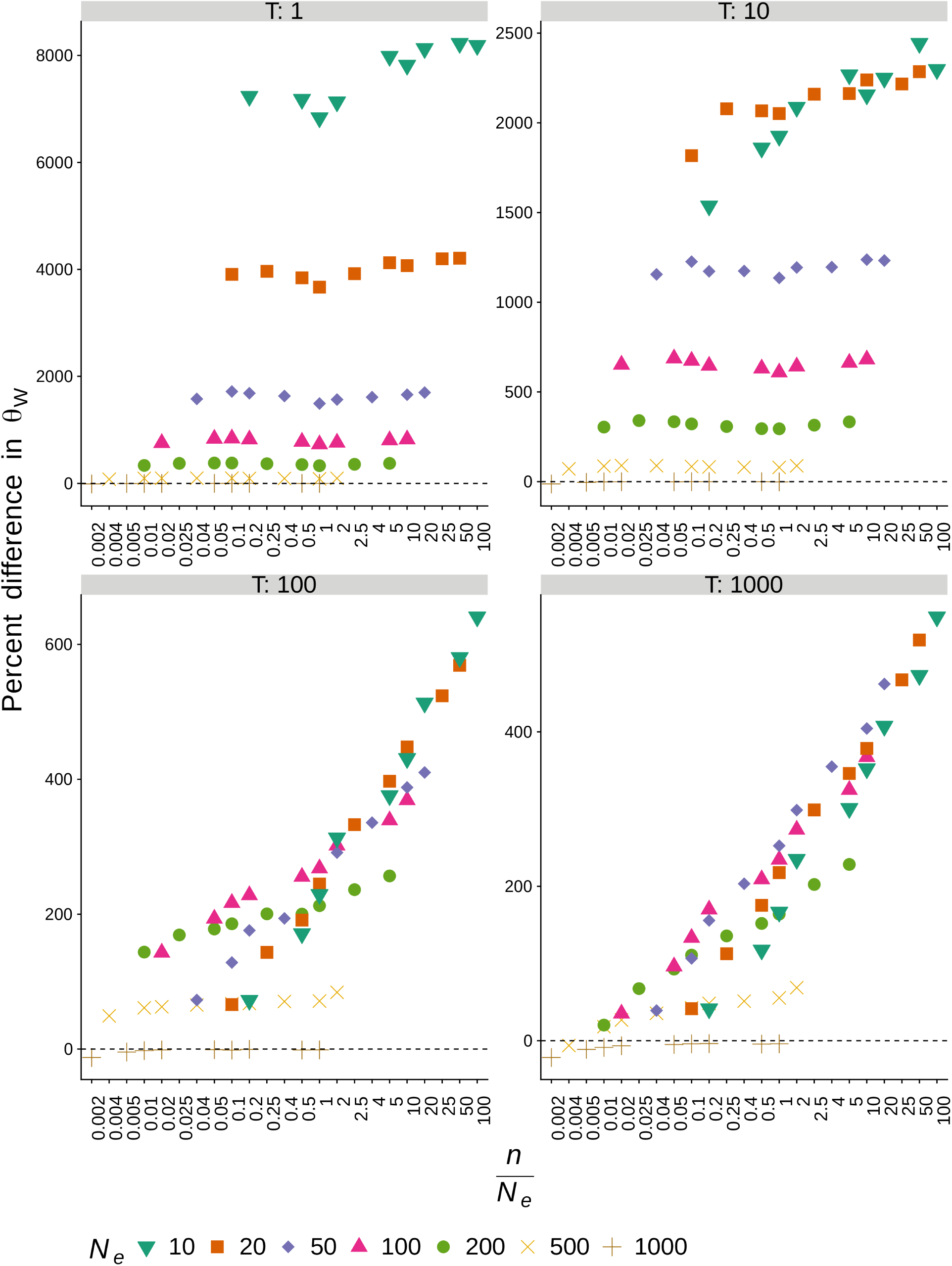
The effect of the relationship between sample size and effective population size (*n/N*_*e*_) on coalescent estimates of *θ*_*W*_ relative to forward calculations of *θ*_*W*_ at different sampling times after a bottleneck (*T* = {1, 10, 100, 1000)}. When the percent difference is positive, the coalescent estimate is larger and thus overestimating *θ*_*W*_, and when the percent difference is negative, the coalescent estimate is smaller and thus underestimating *θ*_*W*_. Note the scales of the y-axes of each panel are different for readability.

As the time since the bottleneck increases, coalescent estimates of *θ*_*W*_ approach forward calculations of *θ*_*W*_. Despite this, there are still statistically significant differences between distributions and means of coalescent and forward *θ*_*W*_s (Kruskal Wallis and Mann-Whitney U tests FDR for most scenarios < 0.001, figure 5). For most scenarios at *T* = 100 and *T* = 1000, coalescent estimates overestimate *θ*_*W*_ (figure 5, figure 8). At these longer times since the bottleneck, the relationship between sample size and effective population size becomes important (figure 7c, d). Since all bottlenecks were from a starting *N*_*e*_ = 1000, these comparisons do not distinguish between percent population size reduction during the bottleneck and final effective population size.

**Figure 8.**
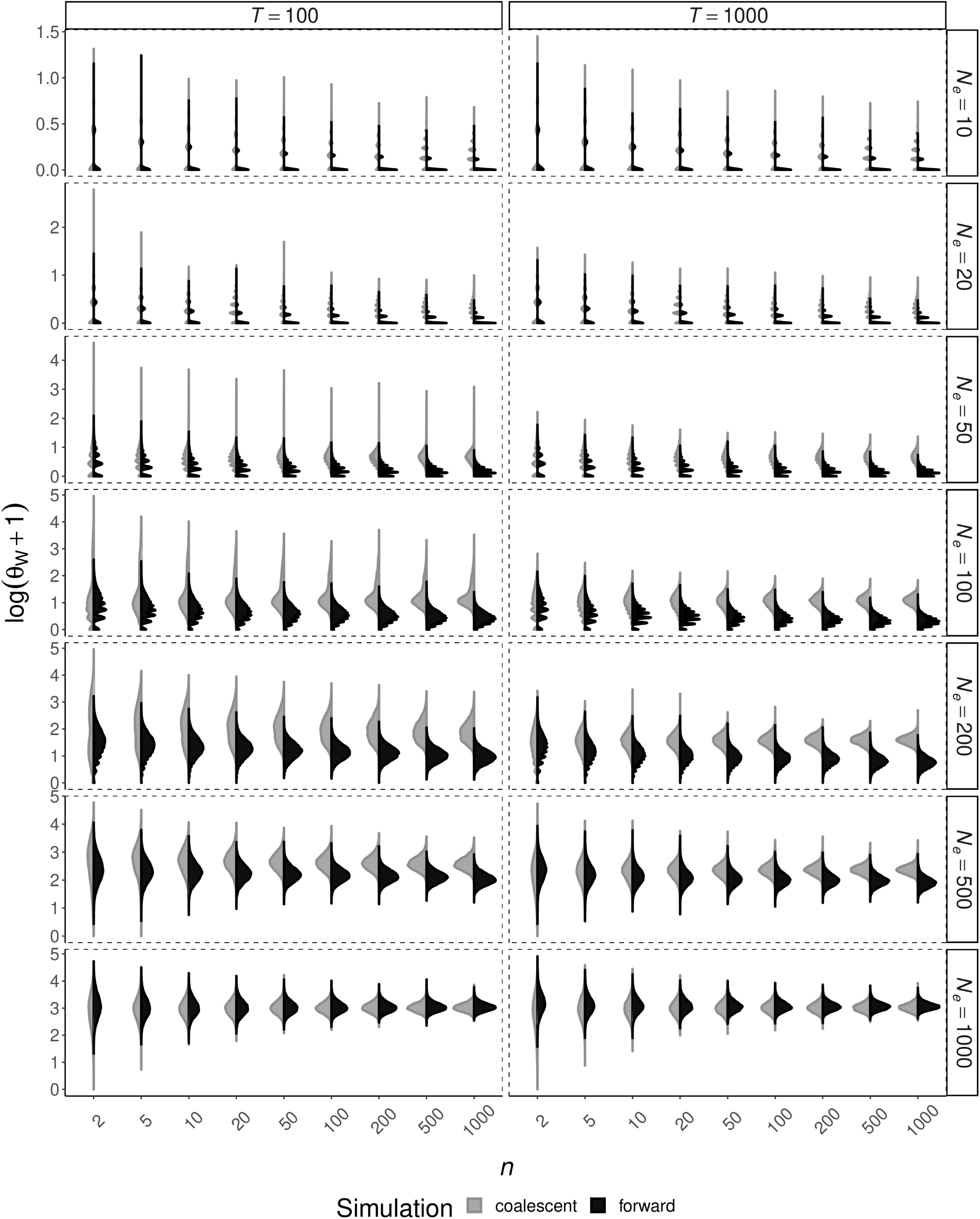
Distributions of *θ*_*W*_ (log transformed for visibility) estimated from coalescent simulations (grey) compared to *θ*_*W*_ calculated from forward simulations (black), across sample size (x axis) and effective population size (vertical facets) for *T* = 100 (left) and *T* = 1000 (right). Note the scales of the y-axes of each panel are different for readability.

## Discussion

To determine if and how standard coalescent models influence estimates of genetic diversity in populations with small effective population sizes, this analysis compares genetic diversity estimates based on coalescent models to genetic diversity calculated in forward simulation ground-truths at a range of effective population sizes, sample sizes, and sampling times since a bottleneck. Coalescent models give unreliable estimates of genetic diversity, as measured by Watterson’s *θ*, regardless of the relationship between sample size and effective population size. This occurs particularly when the population is oversampled with respect to effective population size (sample size exceeds effective population size), and when sampled soon after a bottleneck.

The profound differences between coalescent estimates of *θ*_*W*_ and forward calculations of *θ*_*W*_ soon after a bottleneck show that applying standard coalescent models to bottlenecked populations can give misleading results. Overestimates would have the effect of fitting null coalescent models with incorrectly long times to the most recent common ancestor to estimates of *θ*_*W*_ from real data, thus artificially extending estimates of both bottleneck times and population split times into the past. Underestimates of *θ*_*W*_ that occur after prolonged small effective population sizes would artificially shorten estimates of bottleneck times or population split times.

Bottlenecks are very common, both in species of conservation concern and others, including humans (Hey and Harris 1999). There have been suggestions that standard coalescent models would be inappropriate for bottlenecked populations because a population bottleneck does not merely shorten branches (thus necessitating the rescaling of coalescent time), it increases the likelihood of multiple mergers (Wakeley and Aliacar 2001; Wakeley and Takahashi 2003). However, previous analyses have been restricted to metapopulations, which include migration and extinction/recolonization dynamics (Wakeley and Aliacar 2001), and the results herein show that coalescent models are inappropriate even for the simplest of bottlenecked populations.

The result that coalescent models overestimate *θ*_*W*_ when *n* > *N*_*e*_ is corroborated by previous observations that when *n* > *N*_*e*_ the proportion of singleton polymorphisms increases relative to standard coalescent expectations (Wakeley and Takahashi 2003; Bhaskar, Clark, and Song 2014). Wakeley and Takahashi attribute this excess of singletons to mutations in the generation immediately previous to sampling, at a rate (*θ* * *n*/*N*_*e*_)/2. These mutations account for most of the singletons expected in the entire tree (E[S] = *θ*) (Watterson 1975).

Multiple merger coalescent models, which are more general than the Kingman coalescent, include the **Ξ**-, **Ψ**-, and **Λ**-coalescent-, **Ψ**-, and **Λ**-coalescent -, and **Λ**-coalescent-coalescent (Irwin et al. 2016). These have been developed since the early 2000s, and vary in how they handle multiple merger events as well as the number of descendants per lineage (Árnason and Halldórsdóttir 2015). Though most of the development has so far been theoretical, potential applications to real-world multiple merger scenarios are clear (Schweinsberg 2000; Mohle and Sagitov 2003; Eldon and Wakeley 2005; Birkner and Blath 2008; Tellier and Lemaire 2014; Koskela, Jenkins, and Spano 2015; Montano 2016). At least one software package, Hybrid-Lambda, has been developed to allow the simulation of multiple merger trees, including conversion of the resulting branch lengths to coalescent estimates (Zhu et al. 2015).

When applied to the limpet *Cellana ornata*, which has high reproductive skew, Zhu et al. (2015) showed that multiple merger coalescent models using the Hybrid-Lambda package estimated a population split time of approximately 9,000 generations with a multiple merger model vs. approximately 48,000 generations with the standard Kingman coalescent. Similarly, Árnasson and Halldórsdóttir (2015) applied **Ξ**-, **Ψ**-, and **Λ**-coalescent - and **Λ**-coalescent –coalescent null models (Eldon et al. 2015) to another species, Atlantic cod (*Gadus morhua*), whose high reproductive skew may have contributed to its small effective population size (Hutchinson et al. 2003). They found that previous population divergence time estimates may be too high because of inappropriate applications of the standard coalescent, and that multiple merger models may provide better null hypotheses for testing both split times and natural selection.

Despite the importance of exploring and applying multiple merger coalescent models for species with small effective population sizes and those subject to bottlenecks, a more expedient alternative when simulating null expectations is to use forward instead of coalescent simulations. Forward simulators require more computational resources than coalescent simulators, but effective population sizes of *N*_*e*_ = 1000 and even larger can be manageable, even for tens of thousands of iterations. There are many forward simulation options to choose from, including SFS_CODE (Hernandez 2008), SLiM (Haller and Messer 2017, 2019), and fwdpp (Thornton 2014). SLiM v3 in particular supports non-Wright-Fisher models, which has the potential to extend our understanding of the dynamics of natural populations even further.

Loci from natural populations are typically sampled with ascertainment bias toward polymorphic sites. This would reduce or eliminate sites without polymorphisms, which account for most sites at small effective population sizes, and also accounts for some of the differences in mean *θ*_*W*_ between coalescent and forward simulations. The resulting distributions of both coalescent and forward *θ*_*W*_ would be biased upward, with reduced right skew. This has the potential to reduce apparent differences between coalescent and forward *θ*_*W*_. In addition, *θ*_*W*_ is expected to be more sensitive to the immediate loss of rare variants (and thus segregating sites) caused by a bottleneck than some other commonly used summary statistics such as heterozygosity (Allendorf 1986). Like loci chosen with ascertainment bias, heterozygosity might also reduce the apparent differences between coalescent and forward *θ*_*W*_.

Since natural selection has the effect of reducing effective population sizes at (and near) loci under selection, these results have implications for tests for selection using coalescent-based null models. A similar effect has recently been suggested of the influence of sample size on tests for selection under exponential population growth (Subramanian 2016). The effect of small effective population size and bottlenecks could be easily tested using similar methods as were applied in this study.

This study shows limits in the accuracy of coalescent models with respect to small effective population sizes and bottlenecked populations, including in combination with oversampling. Under these population conditions, standard coalescent models have the potential to produce misleading results and generate incorrect conclusions about population history and structure. In applications to empirical data, it is thus imperative to take into account how robust coalescent models may or may not be to violations of their assumptions.

Thus, standard coalescent models are inappropriate for developing null expectations of genetic diversity for many scenarios with small effective population sizes, especially soon after a bottleneck. This is particularly relevant to studies of species of conservation concern, in which small effective population sizes and bottlenecks are common (Allendorf 1986; Palstra and Ruzzante 2008), but may also be relevant to other species subject to oversampling or bottlenecks. This has implications for studies of species of conservation concern, because coalescent models that profoundly mischaracterize the timings of bottlenecks, population splits, and other demographic events could detrimentally influence management actions. In addition, these mischaracterizations are potentially pernicious in studies of disease population genetics, because repeated bottlenecks and intense selection are classic aspects of disease agent ecology.

## Acknowledgements

Thanks go to Liliana. M. Dávalos and Krishna R. Veeramah for constructive discussions and comments. Simulations were performed on the Stony Brook University Seawulf HPC cluster and Indiana University HPC Mason and Carbonate clusters. This research was supported, in part, by an American Association of University Women Dissertation Fellowship to M. Elise Lauterbur, and NSF-DEB1456455 to Liliana M. Dávalos.

